# Human iPSC-derived pericyte-like cells carrying APP Swedish mutation overproduce beta-amyloid and induce cerebral amyloid angiopathy-like changes

**DOI:** 10.1101/2024.06.07.597867

**Authors:** Ying-Chieh Wu, Šárka Lehtonen, Kalevi Trontti, Riitta Kauppinen, Pinja Kettunen, Ville Leinonen, Markku Laakso, Johanna Kuusisto, Mikko Hiltunen, Iiris Hovatta, Kristine Freude, Hiramani Dhungana, Jari Koistinaho, Taisia Rolova

## Abstract

Alzheimer’s disease (AD) patients often exhibit cerebral amyloid angiopathy (CAA), i.e beta-amyloid (Aβ) accumulation within cerebral blood vessels causing cerebrovascular dysfunction. Pericytes wrap around vascular capillaries, thus regulating cerebral blood flow, angiogenesis, and vessel stability. Vascular dysfunction can promote the development and progression of neurodegenerative diseases, yet the specific contribution of pericytes to AD pathology remains unclear. Here we show that human induced pluripotent stem cell (iPSC)-derived pericyte-like cells (iPLCs) can generate Aβ peptides, and that the cells carrying Swedish mutation in amyloid precursor protein (APPswe) secrete 10 times more Aβ1-42 than the control cells. Additionally, APPswe iPLCs have an impaired capacity to support angiogenesis and barrier integrity, exhibit a prolonged contractile response, and produce increased levels of pro-inflammatory cytokines upon inflammatory stimulation. These functional alterations in APPswe iPLCs are accompanied by transcriptional upregulation of actin cytoskeleton and extracellular matrix organization-related genes. Therefore, the APPswe mutation in iPLCs recapitulates several features of CAA pathology *in vitro*. Our iPSC-based vascular cell model may thus serve as a platform for drug discovery targeting vascular dysfunction in AD.

## Introduction

Alzheimer’s disease (AD) is a progressive neurodegenerative disorder that accounts for 60-70% of dementia cases. Familial AD is caused by gene mutations in the amyloid precursor protein (*APP*) or presenilin-1 and -2 (*PSEN1/2*) genes^1^ that enhance brain deposition of beta-amyloid (Aβ) peptides resulting in amyloid pathology. While most cases of AD are sporadic, investigating familial AD can offer valuable insights, as both familial and sporadic AD share similar pathological features.

Although AD research has mostly focused on neurons, neuronal function is largely dependent on cerebral blood flow (CBF) providing an adequate supply of oxygen and glucose. Recent imaging studies suggest that hypoperfusion is one of the earliest disease-associated changes in the AD brain, promoting cognitive impairment independently from parenchymal amyloid deposition^2–4^. Cerebral amyloid angiopathy (CAA), the accumulation of Aβ in brain vasculature, is considered a fundamental underlying cause of vascular dysfunction in AD, and it is also a risk factor for hemorrhagic stroke^5^. Pericytes and smooth muscle cells (SMCs) are mural cells that wrap around brain capillaries and form arterial vessel walls, respectively. They play an important role in vascular stability and CBF regulation ^6^. CAA promotes the degeneration of endothelial cells, pericytes and SMCs and compromises the integrity of the blood-brain barrier (BBB)^5,7,8^. A single-nucleus RNA-sequencing analysis of AD patient brains showed that AD risk genes are highly expressed in vasculature-associated cell types^9^. Similar findings emerged in the aged mouse brain, where vascular cells exhibited senescent phenotypes or displayed increased expression levels of AD risk genes^10^.

To address the impact of genetic mutations associated with familial AD on pericyte function, we generated pericyte-like cells from three human iPSC lines carrying the KM670/671NL mutation in APP (APPswe), along with seven control lines from healthy individuals (Table 1). Our pericyte-like cells express characteristic pericyte markers, promote a formation of complex tube structure in endothelial cells (ECs) and decrease EC layer permeability, thus demonstrating several normal pericyte functions. These pericyte functions were impaired by the presence of APPswe mutation. Thus, human iPSC-derived vascular cells have the potential to serve as a model for investigating vascular dysfunction in neurodegenerative diseases.

**Table 1.**
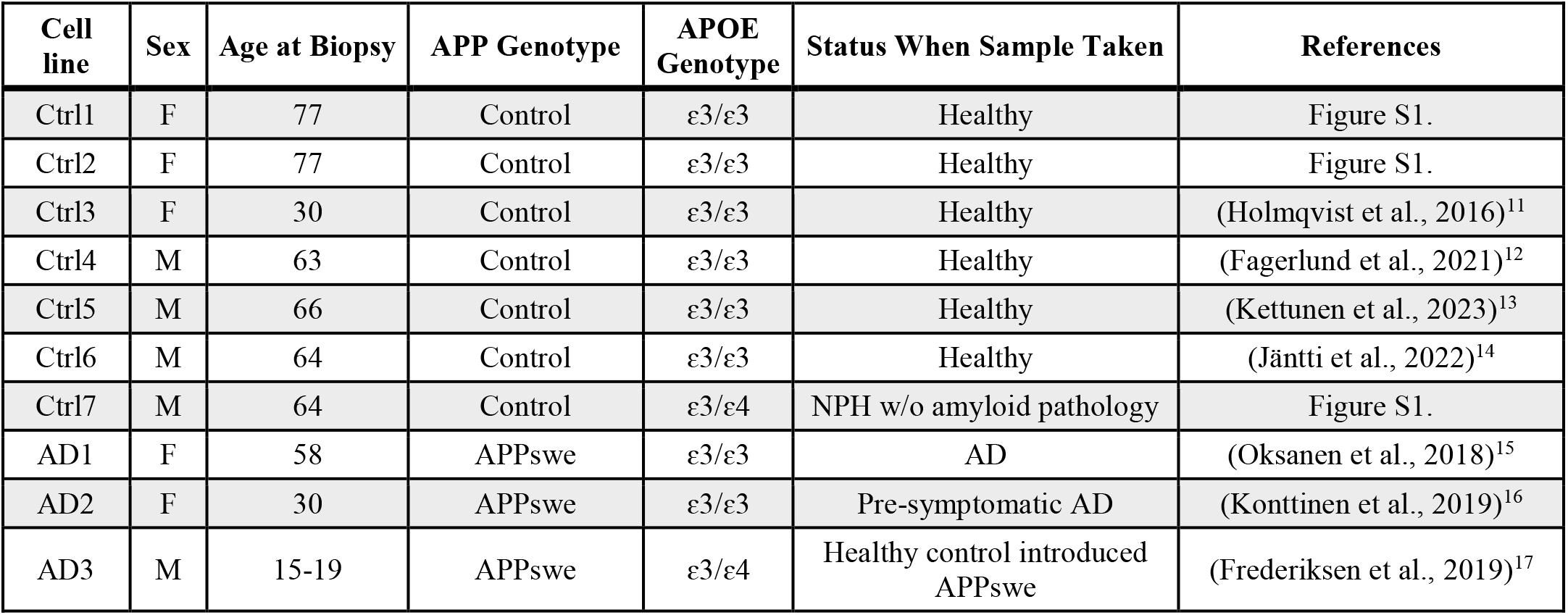
iPSC lines used in this study.

## Results

### iPSCs efficiently differentiate into pericyte-like cells and show similarity to *in vivo* pericytes

Pericyte-like cells were differentiated from human iPSCs (Table 1) via mesodermal route according to a slightly modified protocol from Blanchard et al. 2020^18^. The obtained cells exhibited positive immunoreactivity for mural cell markers platelet-derived growth factor receptor beta (PDGFRβ), neural-glial antigen 2, proteoglycan (NG2), and alpha-smooth muscle actin (α-SMA) (Figure 1 A). Additionally, these cells displayed significantly higher expression levels of mural cell-enriched genes *PDGFRB, DES, LAMA2*, and *PDE7B*, in comparison to undifferentiated iPSCs, iPSC-derived endothelial cells (iECs) and iPSC-derived astrocytes (iAstrocytes) (Figure 1 B). While *DLC1* expression was also elevated in iECs, pericyte-like cells still exhibited a significantly higher expression level than both iPSCs and iAstrocytes. As expected, the cells did not express the pluripotency markers *NANOG, LIN28A*, and *SOX2*, nor the EC marker *CDH5* (Figure S2 A-B). To determine the best timeline to conduct our experiments, we examined the expression levels of pericyte-associated genes in four control lines at days 7, 21, 31, and 50 after the initiation of differentiation. We discovered a gradual rise in the expression of *LAMA2* and *PDE7B* (Figure 1 C), both of which had previously been found to be enriched in pericytes in human post-mortem brain ^9^. The expression of *PDE7B* increased even further on day 50. Additionally, the mural cell marker *CD248* showed a progressive rise until day 31 of the culture period (Figure 1 C). In contrast, the expression of SMC-associated genes *DES* and *ACTA2* decreased significantly from day 7 to day 21 (Figure 1 C). *PDGFRB* and *DLC1* gene expression levels remained stable across the analyzed time points (data not shown). These results imply that the cells acquired a pericyte-like identity between day 21 and 31 post-differentiation, which was selected as the timeframe for subsequent experiments. Given the challenge in definitively distinguishing pericytes from SMCs with the current data, we refer to the generated cells as iPSC-pericyte-like cells (iPLCs) in our study.

**Figure 1.**
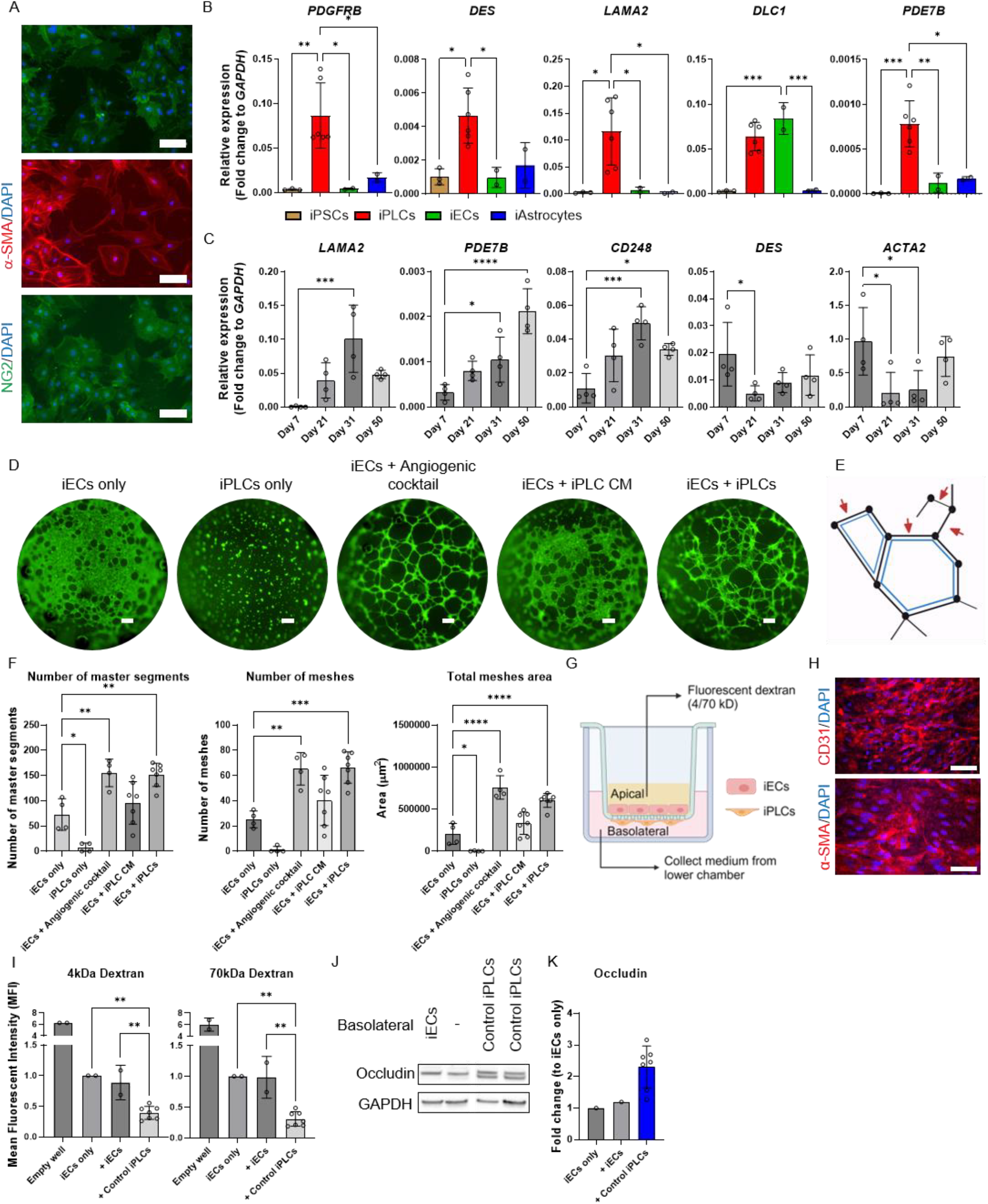
Differentiation and Characterization of iPSC-Derived Pericyte-like cells (i-PLCs). **(A)** Representative immunocytochemistry images of day 21 iPLCs from control lines, stained for PDGFRβ, α-SMA, and NG2. Nuclei are stained with DAPI. Scale bars, 100 μm. The experiments were conducted twice for all lines. **(B)** Relative gene expression levels of *PDGFRB, DES, LAMA2, DLC1*, and *PDE7B* were compared among iPLCs, iECs, iAstrocytes, and iPSCs shown as fold change to *GAPDH*. Statistical differences to iPLCs are indicated (iPLCs, n = 6 lines; iECs, n = 2 lines; iAstrocytes, n = 2 lines; iPSCs, n = 3 lines, experiments were repeated three times for pericytes and two times for rest of cell types). **(C)** Relative gene expression levels of *LAMA2, PDE7B, CD248, DES*, and *ACTA2* were compared across Day 7, 21, 31, and 50 iPLCs shown as fold change to *GAPDH*. Statistical differences to Day 7 iPLCs are indicated (pericytes, n = 4 lines (for all time points); two batches). **(D)** Representative images from 2D tube formation assay showcase various culture combinations: iECs alone, iPLCs alone, iECs exposed to angiogenic cocktail or iPLC CM, and iECs co-cultured with iPLCs. Images were taken 6 hours post-replating. Scale bars, 300 μm. **(E)** The diagram demonstrates master segments (red arrows) and mesh structures (blue polygons). **(F)** Statistical analysis was conducted to compare the number of master segments, meshes count, and meshes area across various culture conditions, with significance determined compared to iECs alone. The experiments were replicated using two batches, each consisting of seven control lines of iPLCs. **(G)** A schematic diagram illustrating the setup for measuring permeability. **(H)** Representative immunocytochemistry images depict iECs and iPLCs on transwell inserts, stained with CD31 and α-SMA, respectively. Nuclei are stained with DAPI. Scale bars, 100 μm. The staining was performed once for two control and two APPswe lines. **(I)** Permeability of fluorescently labeled dextran was evaluated across different culture conditions, with values presented as fold changes relative to iECs-only wells. “+iECs” or “control iPLCs” denotes cells cultured with iECs on the basolateral side of the inserts. Statistical differences compared to iECs only are highlighted. The experiments were replicated in two batches, involving seven control lines of iPLCs. **(J-K)** Western blots were performed for Occludin using samples from one batch of various culture conditions, including iECs alone and co-cultured with iECs or iPLCs on the basolateral side of inserts. GAPDH was used as a loading control, and quantification of the blots was normalized to GAPDH levels. The statistics involving seven control lines of iPLCs. The dots represent average values from batches and technical repeats of each line. Only iECs from (F, I, K) dots represent means of technical repeats within each batch. The data are presented as mean ± SD. Statistical analysis utilized one-way ANOVA with Dunnett’s multiple comparison test, with significance denoted: *p < 0.05, **p < 0.01,***p < 0.001 and ****p < 0.0001.

### iPLCs promote angiogenesis and barrier integrity

The regulation of vascular function and morphology is crucially dependent on the interactions between ECs and vascular mural cells. To explore the angiogenic potential of iPLCs, a 2D tube formation assay was performed. For this purpose, iECs were derived from a line transduced with the transcription factor E26 transformation-specific variant 2 (ETV2). The expression of ETV2 was induced on day three after the initiation of differentiation. Initially, we evaluated the angiogenic capabilities of iECs and iPLCs independently, finding that neither could form clear tube-like structures on a Matrigel layer when cultured alone (Figure 1 D). The analysis was conducted using the ImageJ Angiogenesis Analyzer, which converts images to binary format and identifies tube-like structures. We used parameters such as segment number, mesh number, and area (Figure 1 E) to determine the complexity of the tube structures. Due to converting pixels to binary, even without visible segments or meshes in iECs or iPLCs cultures, the binary images can still produce pseudo structures with low number counts. Subsequently, we examined the ability of iECs to form tubes when subjected to angiogenic factors. Notably, iECs exposed to an angiogenic cocktail (3ng/ml vascular endothelial growth factor (VEGF), 30nM Sphingosine-1-phosphate (S1P), 3ng/ml Phorbol 12-myristate 13-acetate (PMA) and 2.3ng/ml Fibroblast growth factor 2 (bFGF)) successfully formed tubular structures, with significant increases in the numbers of master segments, mesh structures, and total mesh areas (Figure 1 D, F). iPLCs-conditioned medium (CM) did not significantly induce tube formation in iECs. However, direct co-culturing of iECs with iPLCs resulted in the formation of tubular structures comparable to those exposed to angiogenic factors. This was evidenced by significant increases in the numbers of master segments, mesh counts, and mesh areas (Figure 1 D, F). These observations suggest that iPLCs may act as an angiogenic stimulus through direct interactions with ECs.

Additionally, the ability of iPLCs to induce BBB properties in ECs was evaluated using a transwell model. In the experimental setup, iPLCs were placed on the basolateral side of the inserts, while iECs were cultured on the apical side (Figure 1 G). Initially, CD31 and α-SMA stainings were conducted to confirm the attachment of iECs and iPLCs before performing the permeability assay (Figure 1 H). The permeability to 4 kDa and 70 kDa dextran of 7-day-old cocultures was significantly reduced (Figure 1 I) compared to iECs alone and iECs co-cultured on the basolateral side. The Western blot analysis showed the appearance of a lower molecular weight form of the tight junction protein (TJP) Occludin in iEC/iPLCs co-cultures as compared to iECs alone. (Figure 1 J-K). These findings indicate that iPLCs can modulate the formation and integrity of blood vessels, thereby confirming their functional efficacy.

### APPswe iPLCs exhibited differential expression of *in vivo* pericyte markers and expressed α-SMA stress fibers

We expanded our investigation to compare the expression of pericyte genes between iPLCs derived from healthy individuals and those carrying APPswe mutations. Our analysis revealed no significant differences in the expression levels of PDGFRB, LAMA2, DLC1, and CD248 between APPswe and control iPLCs (Figure 2 A). However, a notable reduction in the expression of PDE7B and DES was observed in APPswe iPLCs (Figure 2 B), alongside an increase in ACTA2 expression (Figure 2 B). ACTA2 encodes for α-SMA, a protein associated with the cell contractility. Western blot further suggested an upward trend in α-SMA protein levels in APPswe iPLCs (p = 0.068; Figure 2 C). A detailed analysis of α-SMA staining also revealed distinct organizational patterns. In control iPLCs, α-SMA was diffusely distributed and primarily located at the cell margins, whereas in APPswe iPLCs, there was a noticeable increase in the presence of actin stress fibers (Figure 2 D). The quantification of cells exhibiting actin stress fibers revealed a significant increase in the prevalence of these structures among iPLCs with APPswe mutation (Figure 2 D). These findings suggest potential defects in APPswe iPLCs.

**Figure 2.**
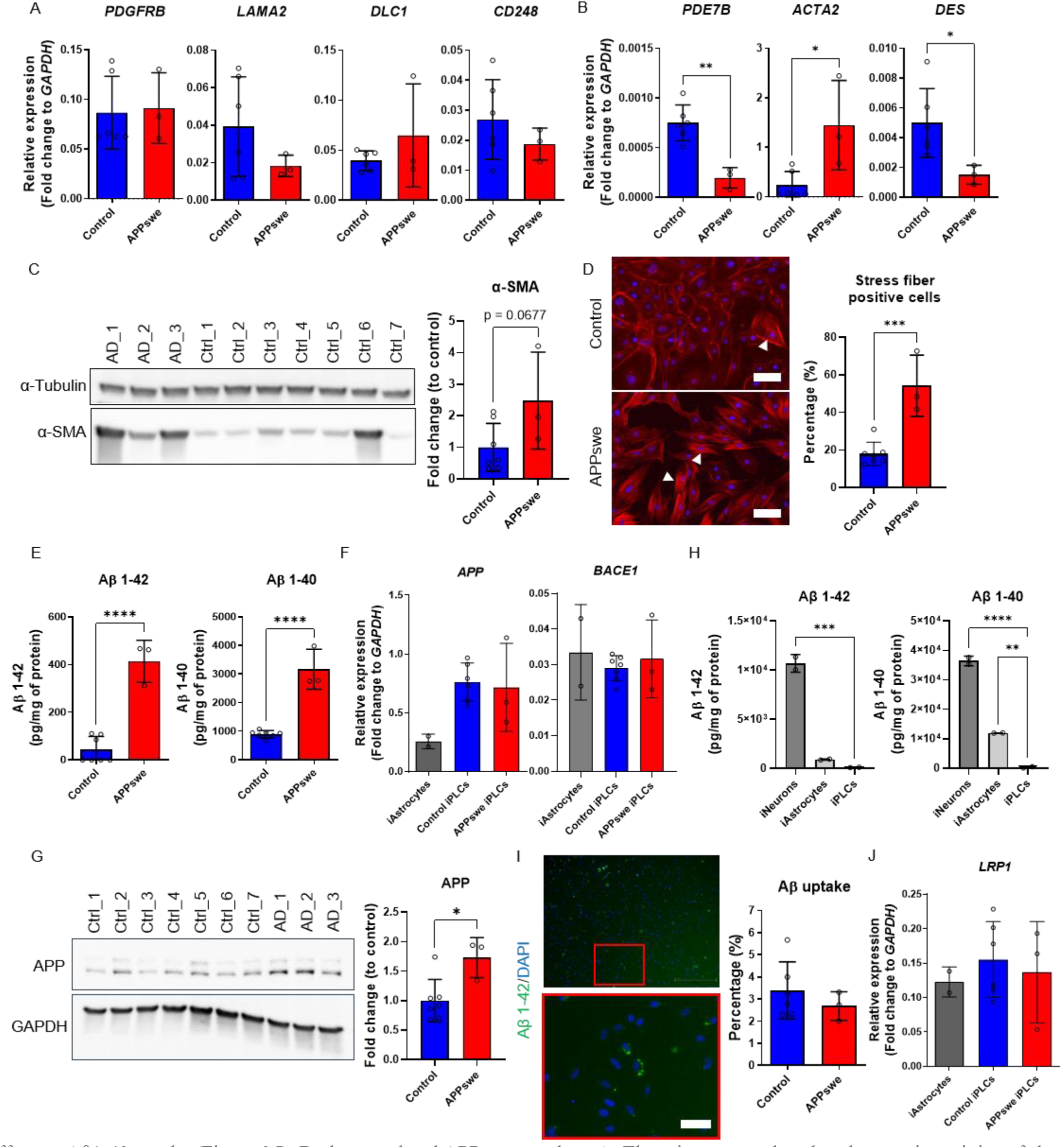
APPswe iPLCs Displayed Altered Expression of Pericyte Markers, α-SMA Stress Fibers, and Amyloid beta Pathology. **(A-B)** Relative expression levels of *PDGFRB, LAMA2, DLC1, CD248* (A), and *PDE7B, ACTA2, DES* (B) were analyzed between control and APPswe iPLCs, normalized to *GAPDH* and presented as fold changes. Statistical differences relative to control are highlighted. This analysis was performed using two experimental batches, including six control and three APPswe lines. **(C)** Representative blots for α-SMA from control and APPswe iPLCs utilized α-tubulin as the loading control. Blot quantification was normalized to α-tubulin levels. This analysis included two experimental batches, with seven control and three APPswe lines. **(D)** Representative images depict α-SMA staining in control versus APPswe iPLCs, with nuclei stained by DAPI. Scale bars, 100 μm. The percentage of cells exhibiting stress fibers was quantitatively compared between control and APPswe cells, utilizing ImageJ’s threshold function to assess cell coverage area. This analysis was conducted over two experimental batches, including seven control and three APPswe lines. **(E)** The levels of Aβ1–42 and Aβ1–40 in the media were measured using ELISA kits, and the obtained values were normalized to the total protein content. Statistical differences compared to control are indicated. The experiments were replicated with two batches, consisting of seven control and three APPswe lines. **(F)** The relative gene expression levels of *APP* and *BACE1* were compared between control and APPswe iPLCs, as well as iAstrocytes, and quantified as fold changes relative to *GAPDH*. The experiments were replicated with two batches, each containing six control lines, three APPswe iPLCs, and two iAstrocyte lines. **(G)** Blot for APP from control and APPswe iPLCs utilized GAPDH as the loading control. Blot quantification was normalized to GAPDH levels. This analysis included one experimental batch, with seven control and three APPswe lines. (H) Aβ1–42 and Aβ1–40 levels in the media were measured for iPSC-neurons, astrocytes, and iPLCs derived from an APPswe individual. The obtained values were normalized to the total protein content. Statistical differences compared to iPLCs cells are highlighted. The experiments were conducted in one batch, consisting of cell types derived from a single APPswe line. **(I)** Images depict iPLCs internalizing HiLyte 488-labeled Aβ1–42. The upper image was captured at 4x magnification, while the lower image was taken at 10x (C). Scale bar = 100 μm. The percentage of cells that internalized Aβ1–42 was quantified from 10x magnification images. The quantification was performed by normalizing the number of engulfed Aβ1– 42-positive cells to the total DAPI counts observed within a single frame. The experiments were replicated with two batches, consisting of six control and three APPswe lines. **(J)** The relative gene expression levels of *LRP1* were compared in control iPLCs, APPswe iPLCs, and iAstrocytes, and were quantified as fold changes relative to *GAPDH*. The experiments were replicated with two batches, consisting of six control, three APPswe, and two iAstrocytes lines. The dots represent the average values obtained from batches and technical repeats of each line. The data are presented as mean ± SD. Statistical analysis was performed using one-way ANOVA with Dunnett’s multiple comparison test (A,B and E) or t-test (D). The significance levels are denoted as follows: *p < 0.05, **p < 0.01,***p < 0.001 and ****p < 0.0001.

### iPLCs can produce and secrete Aβ peptides

The accumulation of Aβ deposits in the brain is a hallmark of AD. Previous research has reported co-labeling of pericytes with Aβ in human AD brain^19,20^. However, it is not clear whether pericytes actively produce Aβ or uptake it from the surrounding extracellular space. Our findings revealed a substantial increase in Aβ1-42 secretion by iPLCs harboring APPswe mutation (413.7 ± 88.4 pg/mg cell lysate over 7 days; Figure 2 E) compared to control cells, where the presence of Aβ1-42 was barely detectable (43.4 ± 54.7 pg/mg cell lysate; Figure 2 E). As expected, iPLCs secreted higher levels of Aβ1-40 than Aβ1-42 (Controls: 894.2 ± 125.8 pg/mg; APPswe: 3165.4 ± 701.7 pg/mg), and the genotype effect was similar to the effect on Aβ1-42 (Figure 2 E). The mRNA levels of *APP* and *BACE1* in both control and APPswe iPLCs were similar to those found in iAstrocytes (Figure 2 F). Notably, APP protein levels were higher in APPswe iPLCs compared to controls (Figure 2 G). Our data suggest that pericytes are capable of producing Aβ and may contribute to the Aβ pathology, although the extent of their contribution is yet to be fully understood. To clarify this, we then compared Aβ production among iPSC-derived neurons (iNeurons), iAstrocytes, and iPLCs to gauge the relative contribution of pericytes to the overall Aβ pathology. The results indicated that APPswe iPLCs secreted on average 100 times lower levels of both Aβ1-42 and Aβ1-40 than APPswe iNeurons when grown at the same cell density for 72 h (Figure 2 H). The findings suggest that the contribution of pericytes to total brain amyloid load in AD is limited.

To investigate Aβ uptake potential of our iPLCs, we introduced fluorescently labeled fibrillar Aβ1-42 to the cultures and incubated them for 24 hours. The quantification of the percentage of cells that internalized Aβ1-42 was performed based on acquired images (Figure 2 I). Only around 3% of control cells ingested Aβ1-42, and the APPswe mutation had no effect on Aβ1-42 uptake (Figure 2 I). Both control and APPswe iPLCs expressed *LRP1* mRNA levels similar to iAstrocytes, which are known to express detectable levels of *LRP1*^21^ (Figure 2 J). Furthermore, when iPLCs were subjected to pHrodo-conjugated zymosan-coated beads, no uptake of these pathogen-mimicking particles was observed (data not shown). Thus, it appears that the phagocytic activity of these iPLCs is low.

which are known to express detectable levels of LRP121 (Figure 2 J). Furthermore, when iPLCs were subjected to pHrodo-conjugated zymosan-coated beads, no uptake of these pathogen-mimicking particles was observed (data not shown). Thus, it appears that the phagocytic activity of these iPLCs is low.

### APPswe mutation significantly altered transcriptome of iPLCs

To further investigate the impact of the APPswe mutation on pericyte-like cells, we conducted transcriptomic analysis on 21-day-old cells. A pairwise comparison identified 687 differentially expressed genes (DEGs) between APPswe and control iPLCs (Figure S3 A, Table S2), with 257 genes upregulated and 430 downregulated. The topmost up- and downregulated genes in APPswe iPLCs are highlighted in red and blue in volcano plot, respectively (Figure 3 A). We used Ingenuity Pathway Analysis (IPA) to identify the pathways most affected by APPswe mutation in iPLCs (Figure 3 B-C, Table S3). Among the five strongest downregulated pathways, we identified RHOGD1 signaling, HIF1α signaling, interleukin (IL)-8 signaling, opioid signaling, and endothelin (ET)1 signaling (Figure 3 B). The ET-1 pathway regulates vasoconstriction^22^ while IL-8 and HIF1α signaling pathways are involved in angiogenesis, inflammation, and metabolic regulation^23,24^. The top five upregulated pathways in APPswe iPLCs were associated with actin/cytoskeleton reorganization, including paxillin signaling, actin cytoskeleton signaling, agrin interactions at neuromuscular junction, signaling by RHO family GTPases, and PAK signaling (Figure 3 C). Among the DEGs shared by these pathways there was a significant number of genes related to myosin chains, smooth muscle actin, and integrins (Figure 3 D). Another pathway enrichment analysis using Pathview revealed significant changes in vascular smooth muscle contraction and cardiomyopathy-related pathways, extracellular-receptor interaction and focal adhesion pathways. Detailed examination of genes within these pathways identified *ITGA2, ITGA4, ITGA6*, and *MYLK*, which are involved in cytoskeleton reorganization as described in IPA analysis (Figure 3 F, Table S4). Gene ontology analysis further underscored an enrichment in biological processes associated with vascular and muscle functions and extracellular matrix/structure organization (Figure 3 G, Table S5-7), which are crucial for vascular stability and pericyte fate determination^25^. This analysis also emphasized an enrichment in cellular components of contractile fibers and myofibers (Figure 3 G, Table S5-7). In summary, our transcriptomics data suggests that APPswe iPLCs may exhibit altered functionalities such as cell contractility, inflammatory response, and metabolism regulation.

**Figure 3.**
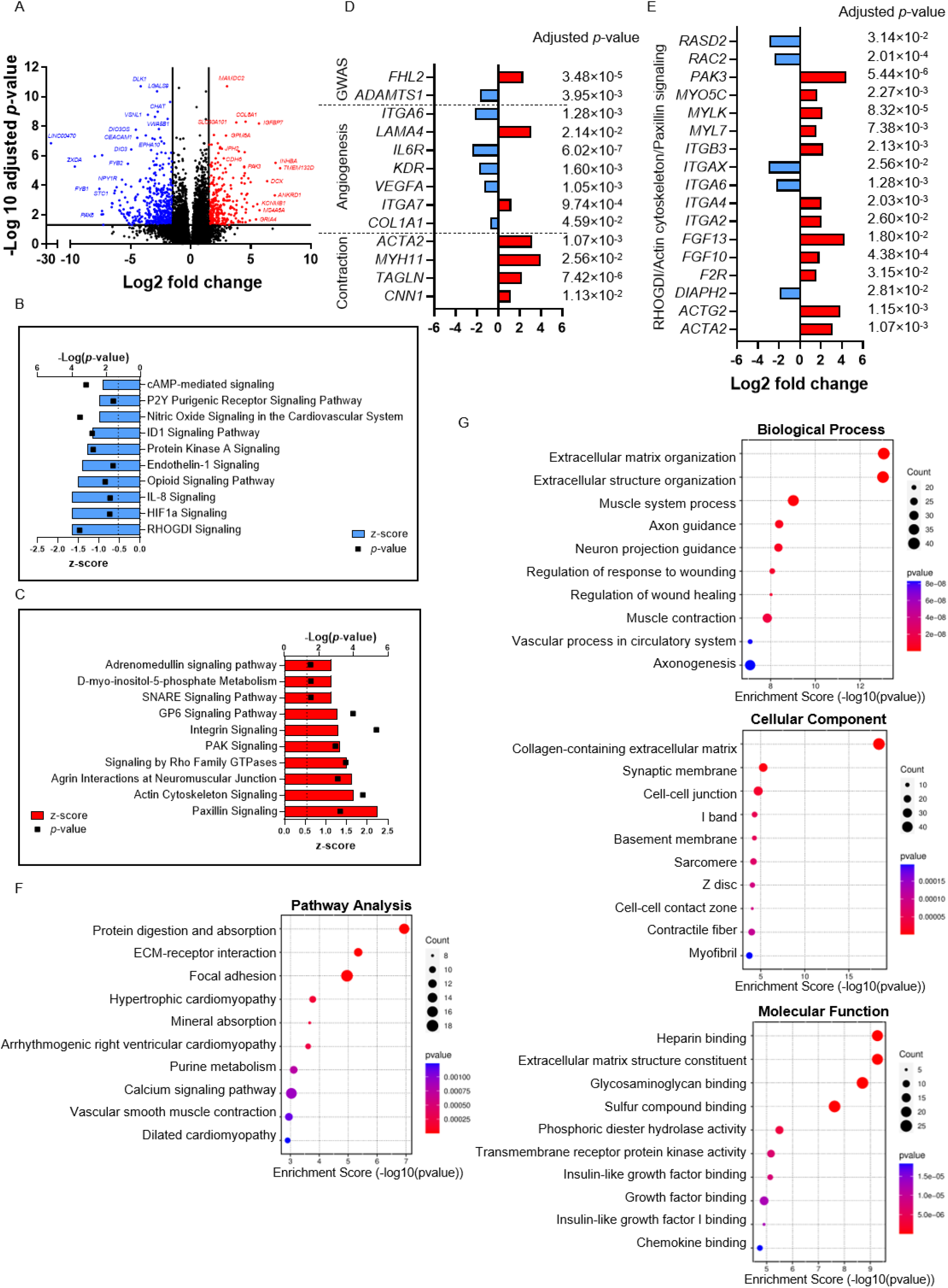
APPswe iPLCs Exhibit an Altered Transcriptome. **(A)** Volcano plot depicting DEGs between control and APPswe iPLCs (cutoffs: Adjusted p-value <0.05 and absolute log2 fold change >1.5). The analysis included seven control and three APPswe lines. **(B-C)** Ingenuity Pathway Analysis to identify the top 10 downregulated (C) and upregulated (D) canonical pathways with the highest z-scores (p-value < 0.05) when comparing control pericytes to APPswe pericytes. **(D)** Genes related to gene risk associated with AD from GWAS, angiogenesis and pericyte contraction process. (cutoffs: Adjusted p-value <0.05 and absolute log2 fold change >1). **(E)** Genes involved in RHOGDI, Paxillin and Actin Cytoskeleton signaling pathways. **(F)** Pathview pathway analysis to identify the top 10 affected pathways with the highest enrichment score when comparing APPswe pericytes to control pericytes. **(G)** Gene ontology (GO) enrichment analysis revealed pathways enriched in Biological Process, Molecular Function, and Cellular Component in APPswe pericyte-like cells relative to controls.

### APPswe iPLCs produce higher levels of MCP-1 after inflammatory stimulation

Pericytes can sense inflammatory stimuli and activate innate immune responses, such as the release of pro-inflammatory cytokines and overexpression of adhesion molecules, including intercellular adhesion molecule (ICAM)-1 and vascular cell adhesion molecule (VCAM)-1^26,27^. Given that our transcriptome analysis (Figure 3 D) showed a downregulation of IL-8 signaling pathway in APPswe iPLCs, we tested the effect of a 24-hour exposure to a combination of proinflammatory cytokines, tumor necrosis factor-alpha (TNFα) and IL-1β (Figure 4 A) on these cells. Following the exposure, inflammatory mediators, IL-6 (two-way ANOVA, *p* = 0.0427), IL-8 (*p* = 0.0028), monocyte chemoattractant protein (MCP)-1 (CCL2) (*p* < 0.0001), regulated on activation, normal T cell expressed and secreted (RANTES; CCL5) (*p* = 0.0267), and soluble VCAM-1 (*p* = 0.0156), were secreted by iPLCs. It is worth noting that bacterial lipopolysaccharide (LPS) failed to induce this response (data not shown), potentially due to an extremely low expression levels of Toll-like receptor (*TLR) 2* and *TLR4* genes (data not shown). Upon stimulation with TNFα and IL-1β, APPswe iPLCs secreted significantly higher levels of MCP-1 and showed a rising trend in soluble VCAM-1 (p= 0.0517) compared to control cells (Figure 4 A). No significant genotype effect was observed on the levels of the remaining inflammatory mediators tested. This suggests that APPswe iPLCs are sensitive to inflammatory stimuli, resulting in an increased release of certain pro-inflammatory mediators.

**Figure 4.**
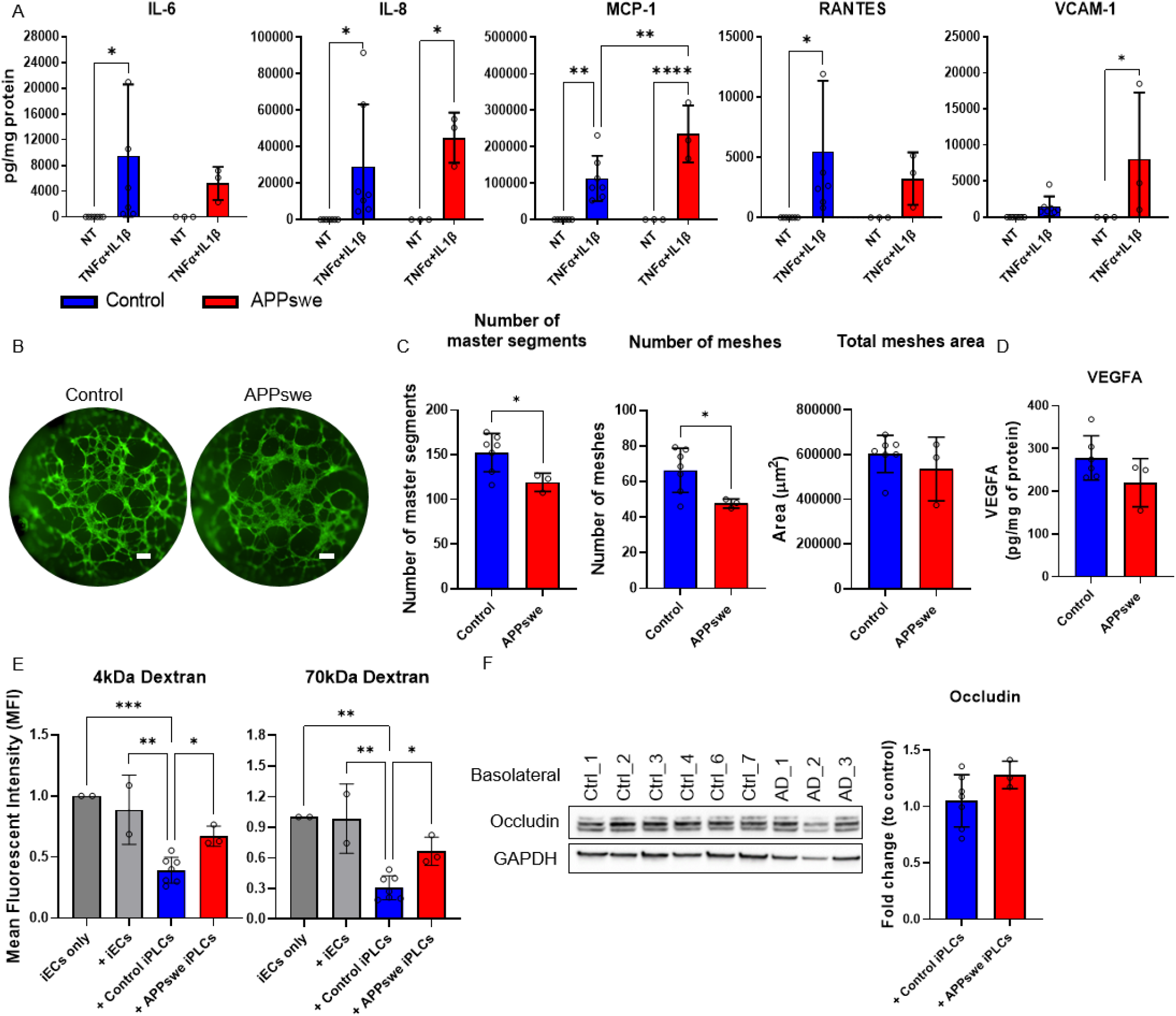
APPswe iPLCs display functional impairments in their response to inflammation, assisting angiogenesis, and maintenance of barrier integrity. **(A)** IL-6, IL-8, MCP-1, RANTES, and VCAM-1 concentrations were measured in iPLCs culture media after 24 hours of TNFα and IL-1β stimulation. Results were normalized to total protein content. Statistical differences between treatment and genotype were indicated. The experiments were replicated with two batches, involving seven control and three APPswe lines. **(B-C)** 2D tube formation assays involved culturing iECs alongside both control and APPswe iPLCs (B), scale bars, 300 μm. Statistical comparisons were made regarding the number of master segments, number of meshes, and mesh area between co-cultures of control and APPswe iPLCs with iECs (C), highlighting significant differences compared to controls. This analysis was conducted across two experimental batches, including seven control and three APPswe lines. **(D)** VEGFA levels in lysates from control and APPswe iPLCs were measured and normalized to total protein content. Statistical differences relative to controls are indicated. This analysis included two experimental batches, with six control and three APPswe lines. **(E)** Permeability to fluorescently labeled dextran was assessed in cultures of iECs alone, iECs co-cultured with iECs, control or APPswe iPLCs on the basolateral side of inserts, using 4 and 70 kDa fluorophore-conjugated dextran. Statistical differences relative to iECs co-cultured with control iPLCs are noted. This evaluation was repeated across two experimental batches, including seven control and three APPswe lines. **(F)** Representative blots of Occludin from control and APPswe iPLCs used GAPDH as the loading control, with quantification normalized to GAPDH levels. The analysis including two experimental batches with seven control and three APPswe lines. The dots represent the average values obtained from batches and technical repeats of each line. The data are presented as mean ± SD. Statistical analysis was performed using two-way ANOVA with Bonferroni multiple comparison test (A, F and G) or t-test (C, D). The significance levels are denoted: *p < 0.05, **p < 0.01,***p < 0.001 and ****p < 0.0001.

### APPswe iPLCs impair angiogenesis and barrier integrity

Given the observed changes in angiogenesis-related gene expression within the APPswe iPLCs transcriptome compared to controls (Figure 3 B), we used tube formation and dextran permeability assays to evaluate the ability of APPswe iPLCs to support normal EC functions. In co-cultures of iECs with APPswe iPLCs, the network of self-assembled tubes was less complex compared to those formed with control iPLCs (Figure 4 B). Statistical analysis revealed a significant reduction in the number of segments and meshes in iECs co-cultured with APPswe iPLCs relative to control cells, although no differences were observed in total mesh area (Figure 4 C). These results suggest that APPswe iPLCs have a diminished capacity to support angiogenesis. We then checked the VEGFA levels by ELISA due to significantly reduced transcript levels in APPswe iPLCs compared to controls (Figure 3 B). However, no difference was observed in VEGFA levels within cell lysates between APPswe and control iPLCs (Figure 4 D), and the secretion of soluble VEGFA into the culture medium was undetectable for both groups (data not shown).

Consistent with previous findings, the permeability of iECs to both 4 kDa and 70 kDa fluorescently labeled dextran was notably reduced in co-cultures with control iPLCs, as opposed to iEC-only cultures and co-cultures with iECs on the basolateral side (Figure 4 E). However, permeability significantly increased when iECs were co-cultured with APPswe iPLCs compared to those with control cells, aligning with iEC-only cultures and those co-cultured with iECs on basolateral side (Figure 4 E). This observation indicates that APPswe iPLCs potentially undermine the integrity of the endothelial barrier. Despite these changes, no significant differences in Occludin levels were detected between the groups (Figure 4 F), indicating that further research is needed to understand the underlying mechanisms. Overall, these results demonstrated that the APPswe mutation decreases the complexity of vascular-like network and compromises the integrity of EC barrier *in vitro*.

### APPswe iPLCs exhibit a prolonged response to ET-1 treatment

Since we had detected a higher prevalence of stress fibers (Figure 2 D) and an upregulation of cytoskeleton reorganization pathways in APPswe iPLCs (Figure 3 E-G), we aimed to further investigate potential defects in pericyte contractility. To validate the contractile ability of iPLCs in response to vasoconstricting and vasodilating signals, we exposed our cultures to ET-1 or adenosine triphosphate (ATP) respectively. We then monitored cell contraction process using the xCELLigence system.

This system measures the electric impedance of cell layer converting it into a cell index that reflects changes in the cell surface area. The iPLCs exhibited immediate contraction in response to ET-1 administration, as shown by a reduction in cell index (Figure 5 A). After the initial contraction, there was an increase in cell index 20-30 minutes post-ET-1 treatment, suggesting cell relaxation. In contrast, when ATP was used as a vasodilator, it caused an elevation in the cell index (Figure 5 A), indicating a potential relaxation of the cells. Further analysis of the contraction/relaxation dynamics, as inferred from the slope changes, showed no significant differences between cells exposed to 10 nM and 100 nM of ET-1 (Figure 5 B). Consequently, we chose 10 nM ET-1 for subsequent experiments.

**Figure 5.**
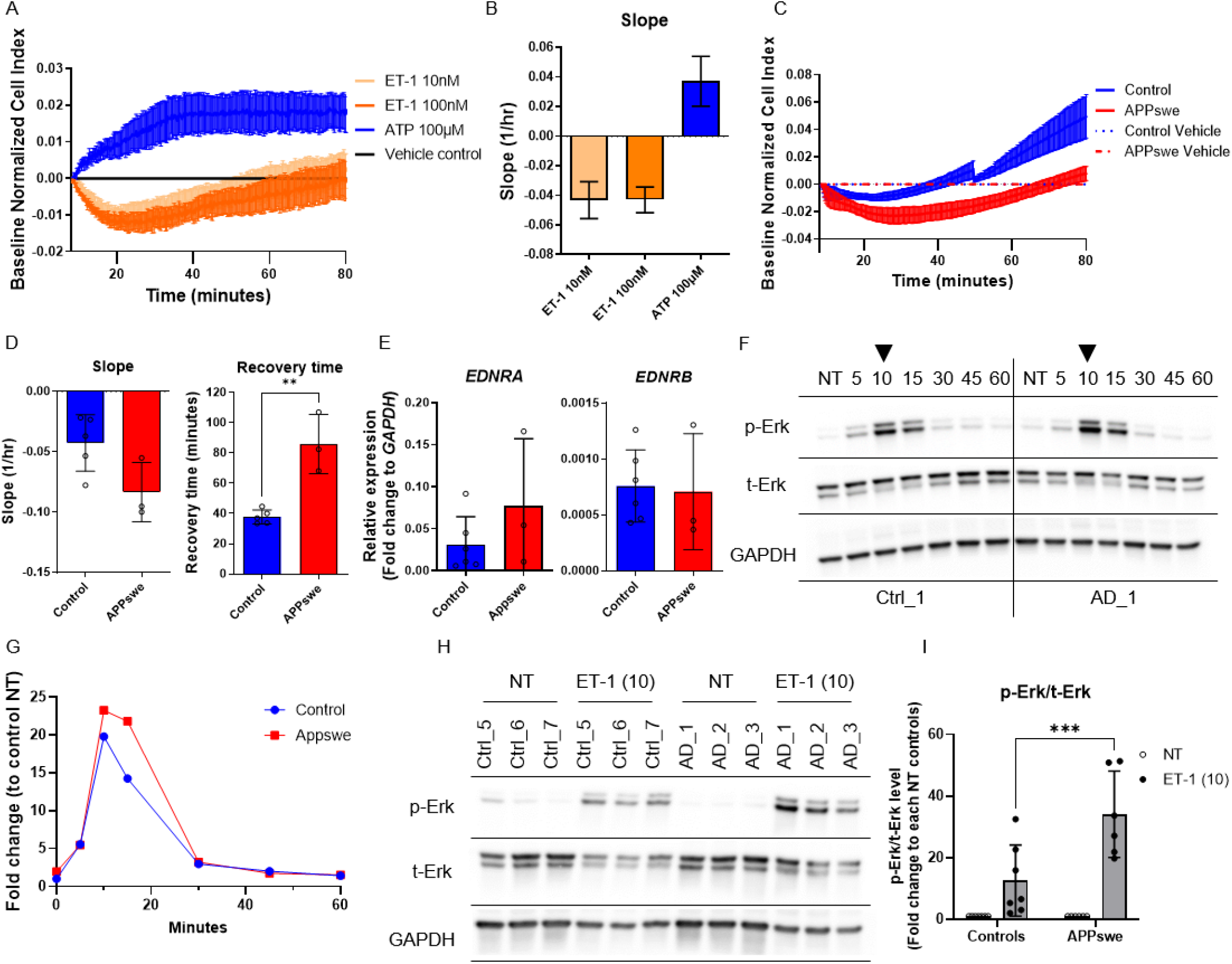
APPswe iPLCs exhibited a hypercontractile phenotype. **(A-B)** Electrical impedance measurements were employed to evaluate the contractile response of iPLCs to various concentrations of ET-1 and ATP. This analysis was conducted in two experimental batches using two control lines. Response curves were normalized to vehicle control, producing a cell index that depicted the response (A), while the response slope, indicating contraction speed post treatment, was also presented (B). **(C)** The cell index of the contractile response of control and APPswe iPLCs to ET-1 treatment, across three experimental batches involving five control and three APPswe lines. **(D)** Statistical analysis compared contraction strength and recovery time to normal size, with significance indicated compared to the controls. The experiments were replicated with three batches, involving five control and three APPswe lines. **(E)** Gene expression levels of *EDNRA* and *EDNRB* were compared between control and APPswe iPLCs, quantified as fold changes relative to GAPDH. The experiments were replicated with two batches, involving six control and three APPswe lines. **(F-G)** Blots for phosphorylated (p)-Erk and total (t)-Erk from control and APPswe iPLCs were conducted at time intervals of 5, 10, 15, 30, 45, and 60 minutes post-ET-1 treatment, as well as for non-treated (NT) cells, using GAPDH as the loading control (F). Quantification was normalized to GAPDH levels, with each time point further normalized to the NT well of the control line (G). This analysis was carried out one experimental batch, involving one control and one APPswe line. **(H-I)** Blots for p-Erk and t-Erk from control and APPswe iPLCs were obtained 10 minutes after ET-1 treatment and from NT cells, with GAPDH serving as the loading control (H). Blot quantification was normalized to GAPDH levels, and treatment groups were further normalized to the NT wells of their respective lines (G). This analysis was performed in a single experimental batch, including seven control and three APPswe lines. The dots represent the average values obtained from batches and technical repeats of each line. The data are presented as mean ± SD. Statistical analysis was performed using two-way ANOVA with Bonferroni multiple comparison test (A, F and G) or t-test (C, D). The significance levels are denoted: *p < 0.05, **p < 0.01,***p < 0.001 and ****p < 0.0001.

After confirming basic contractile responses, we investigated the behavior of both control and APPswe iPLCs. In comparison to the control group, APPswe iPLCs tended to have a steeper contraction slope (p=0.0577) and required a significantly prolonged recovery period to attain normal cell index (Figure 5 C-D). Next, we assessed the transcriptomics data to determine the expression levels of ET-1 receptors. The iPLCs expressed both EDNRA and EDNRB genes but the former was expressed on average 10-times higher than the latter (data not shown). This suggests that endothelin receptor type A may be the primary ET-1 receptor in iPLCs. Nonetheless, there were no significant differences in the expression of these genes between the genotypes. These results were confirmed using RT-qPCR (Figure 5 E). Given that ET-1 receptors are G-protein-coupled receptors (GPCRs) and that Erk is a common downstream pathway, we collected cells at various time points following ET-1 treatment to observe alterations in downstream signaling. The levels of Erk phosphorylated at Thr202 and Tyr 204 (p-Erk) peaked in both control and APPswe iPLCs around 10 minutes post-treatment, with a decline thereafter (Figure 5 F). Interestingly, the p-Erk/t-Erk ratio at 10 minutes post-ET-1 treatment increased more in APPswe iPLCs than in controls (Figure 5 H-I). These observations suggest that APPswe iPLCs exhibit a hypercontractile phenotype with prolonged recovery post-ET-1 exposure. However, further research is needed to confirm the significance of the elevated p-Erk/t-Erk ratio.

## Discussion

In this study, we generated human iPLCs carrying genetic variants linked to familial AD. These iPLCs expressed high levels of pericyte marker genes. Interestingly, some pericyte marker genes such as *PDE7B, ACTA2*, and *DES* exhibited differential expression in APPswe iPLCs compared to controls. These changes are likely due to the intrinsic properties of the APPswe mutation in iPLCs rather than differentiation guidance failures.

A key discovery of our study is a novel mechanism potentially contributing to CAA pathology. Previous studies have reported Aβ production by cerebrovascular cells^29,30,^. However, the significance of these findings has remained ambiguous because Aβ accumulation in the vasculature is generally thought to originates predominantly from neurons^30^. Indeed, we see that iPLCs produce substantially less Aβ than neurons. However, our findings reveal that APPswe iPLCs produced on average 10 times higher levels of Aβ1-42 and 3.5-fold higher levels of Aβ1-40 as compared to controls. Given their local environment, pericytes could still contribute significantly to local high Aβ concentrations, potentially affecting nearby vascular cells.

We also showed that APPswe iPLCs have an impaired capacity to support angiogenesis and barrier integrity, exhibit a prolonged contractile response, and produce increased levels of pro-inflammatory cytokines upon inflammation.

The hypercontractile phenotype of APPswe iPLCs was accompanied by an increased mRNA expression of *ACTA2*, which encodes for α-SMA, and other cytoskeleton-regulating proteins. *ACTA2* and *DES* are crucial for cytoskeleton organization and structural maintenance. The contractile functions in SMCs are mostly dependent on contractile proteins such as α-SMA, smooth muscle myosin heavy chains, and calponin^31^. While it’s unclear whether pericytes utilize the same contraction mechanisms as SMCs, they are thought to modulate CBF through contraction. Our data also revealed an increase in the prevalence of stress fibers in APPswe iPLCs. Stress fibers are linked with contractile activity in myofibroblasts and cardiomyocytes^32^. Increased expression of contractile proteins has been reported in SMCs obtained from AD individuals^33^. Moreover, Yilmaz-Ozcan and coworkers have demonstrated that the small interfering RNA-induced decrease in α-SMA expression levels resulted in a reduction in pericyte contraction in mouse brain^34^. These findings further provide support for the hypothesis that there is a correlation between pericyte contraction and α-SMA expression. Previous publications have also shown that Erk and MAPK pathways can regulate myosin light chain phosphatase, thereby contributing to sustained smooth muscle contraction^35,36^. Our examination of Erk phosphorylation following exposure to vasoconstrictor ET-1 revealed enhanced Erk pathway activation in APPswe iPLCs (Figure 5 F-I). This suggests that the hypercontractility phenotype, characterized by a steeper slope and prolonged recovery time, may stem from amplified Erk signaling, although additional evidence is required to firmly establish this conclusion.

The hypercontractile phenotype that we observed in APPswe iPLCs may also be linked to Aβ pathology. A previous study found that the accumulation of Aβ has a constricting effect on capillaries in patients with cognitive decline. This was achieved by eliciting the ET-1 signaling pathway in pericytes^37^. Similarly, *in vitro* human pericytes exhibit compromised contraction and relaxation after the exposure to exogenous Aβ^38,39^. Thus, the hypercontractile phenotype observed in APPswe iPLCs from our model is consistent with prior studies and could potentially be associated with decreased in CBF in patients with AD. In conclusion, our results demonstrate a hypercontractility phenotype in AD iPLCs, potentially connected to defects in CBF regulation *in vivo*. This phenotype correlates with elevated expression levels of contractile proteins, amplified Erk signaling, and Aβ pathology. Further research is required to fully understand the underlying mechanisms.

In dextran permeability studies, APPswe iPLCs exhibited a compromised ability to enhance the tightness of EC layer. This effect could result in BBB leakage, commonly seen in CAA patients, potentially leading to neuroinflammation and accelerating the progression of AD^40^. While TJPs were upregulated in ECs co-cultured with control iPLCs compared to ECs alone, there was no difference between ECs co-cultured with APPswe iPLCs and controls. This suggests that the effect could be due to variations in iPLCs attachment or the involvement of other TJPs or molecules affecting permeability. Moreover, APPswe iPLCs secreted significantly higher levels of the chemokine MCP1. It is known that overexpression of MCP-1 and VCAM-1 in the endothelium causes monocytes to adhere strongly to the endothelial layer^41,42^. This phenomenon has been linked with endothelial barrier dysfunction^43,44^. Thus, our *in vitro* models containing APPswe iPLCs successfully replicate some features of CAA.

Angiogenesis plays a crucial role in the development and adulthood. Impaired angiogenesis can decrease vessel density, reduce CBF and nutrient transport, and thus potentially worsen the progression of AD. Previous studies indicated that reduced capillary density may result from both vessel degeneration and insufficient angiogenesis due to a lack of angiogenic factors. Our tube formation assay showed that iECs co-cultured with APPswe iPLCs formed slightly less complex tube structures compared to those with control cells. We identified angiogenesis-related genes that were significantly reduced in APPswe iPLCs (Figure 3 B), potentially linking them to the observed phenotype. *VEGFA*, which mural cells can produce to promote vessel formation, along with its receptor, *KDR*^45,46^, were significantly reduced in APPswe cells at the mRNA level. However, no differences were observed at the protein level in cell lysates, and the soluble form of VEGFA was undetectable in the medium, likely due to technical limitations. *IL6R* was also downregulated at the transcription level in APPswe iPLCs. IL-6/IL6R signaling is known to regulate EC proliferation and migration, influencing angiogenesis^47,48^. Additionally, collagen I (*COL1A1*), which supports and guides endothelial cell migration^49^, was reduced in APPswe iPLCs. It has been implicated in angiogenesis-related pathway changes in diabetic retinopathy^50^. Similarly, *ITGA6* was reduced in APPswe cells. It is involved in endothelial morphogenesis by regulating CXCR4 expression to protect established endothelial tubes^51^, similarly showed reduced expression in APPswe cells. Overall, our study highlights transcriptional and functional defects in the support of angiogenesis by APPswe iPLCs and identifies potential molecular targets for further research.

In conclusion, our study shows that APPswe iPLCs induce CAA-like changes, leading to increased BBB permeability and defective angiogenesis and vasoconstriction *in vitro*. Utilizing human iPSC-derived models of brain vascular cells may enhance our understanding of various diseases, including CAA, and help identify key pathways and the development of novel therapies. Given the high prevalence of vascular dysfunction in AD, combination drugs targeting both Aβ pathology and vascular dysfunction could be a promising therapeutic strategy.

## Acknowledgements

We express our gratitude to Anne Nyberg, Agnes Viherä, and Erja Huttu for their valuable assistance in the characterization of iPSCs. We extend our appreciation to Dr. Kristine Freude for generously providing the BIONi010-C Swedish line for our study. Also, we express our gratitude to Dr. Vesa Olkkonen for generously lending us the xCELLigence device, which we utilized to measure pericyte contraction. Additionally, we acknowledge the Biomedicum Functional Genomics Unit (FuGu) at the University of Helsinki for their support in providing bulk RNA-seq services. We extend our gratitude to the Genome Biology Unit and the Biomedicum Virus Core of the GoEditStem platform, as well as Biocenter Finland, for their support in providing plasmid and virus packaging services. This project has received funding from the European Union’s Horizon 2020 research and innovation program under the Marie Skłodowska-Curie grant agreement No. 813294 (J.K. and Y.C.W.). The Sigrid Juselius Foundation (J.K.,Š.L. and MH), The Academy of Finland (grant 334525, J.K., grant 338182, MH), and the Jane and Aatos Erkko Foundation (Š.L.). The funders had no role in the study design, data collection, or interpretation. We declare no competing interests.

## Author contributions

Y.C.W. conducted the majority of experiments, analyzed the data, and wrote the initial draft of the manuscript. K.T. and I.H. performed the normalization of bulk RNA-seq data and conducted the analysis of DEGs. R.K. and H.D. generated doxycycline inducible ETV2 iPSC line. P.K. provided an NGN2 iPSC line. K.F. provided a Swedish mutation iPSC line for the study. T.R. conceptualized the project, provided supervision, and contributed to manuscript revision and comments. Š.L. and J.K. provided valuable input on the manuscript and supervised the project. V.L., M.L., J.Ku. provided patient fibroblasts for the study. All authors have thoroughly reviewed and approved the final version of the manuscript for publication.

## Data availability

Any data and materials available for sharing will be provided under a Material Transfer Agreement. The raw sequencing data and metadata are accessible on the EU Open Research Repository (Pilot) in Zenodo, identified by 10.5281/zenodo.11488682 (https://doi.org/10.5281/zenodo.11488682). Please note that the raw RNA-seq data is kept strictly confidential to uphold patient privacy and confidentiality.

## Materials and Methods

### Patients and iPSCs

AD iPSC lines were generated from two individuals harboring the APPswe mutation; AD1 was diagnosed with AD, while AD2 was pre-symptomatic without a clinical diagnosis. To confirm the causality of the APPswe mutation and AD phenotype, AD3 was kindly provided by Dr. Kristine Freude’s lab, in which KM670/671NL mutations have been introduced in iPSCs from a healthy individual by using CRISPR-Cas9^17^. Seven controls from healthy male and female individuals were used in this study. To reduce the complexity of the genetic background and investigate the phenotype developing from the APPswe mutation, the selected controls (Ctrl1-6) and APPswe mutation (AD1-2) individuals are all carrying apolipoprotein (APOE) ε3/ε3, the AD risk-neutral allele. However, AD3 was carrying ε3/ε4 as its parental line. Therefore, we introduced Ctrl7, which carries APOE ε3/ε4 allele and comes from a patient diagnosed with normal pressure hydrocephalus (NPH) without amyloid pathology, as an additional control for AD3 (Table 1). All iPSC lines have been tested for pluripotency markers, karyotyped, and shown to be capable of forming embryoid bodies and differentiating into three germ layers.

### Generation and maintenance of iPSCs

Dermal biopsies were collected for the generation of iPSCs. All samples were collected with informed consent and approval from the committee on Research Ethics of Northern Savo Hospital District (license no. 123/2016). Fibroblasts were isolated and enriched following the previous publication by Korhonen et al., 2015^52^. Then, somatic cells were reprogrammed to iPSCs with CytoTune-iPS 2.0 Sendai Reprogramming Kits (Thermo Fisher Scientific) as described in the previous publication by Holmqvist et al., 2016^11^. For Ctrl1 and Ctrl2 lines: fibroblasts were reprogrammed at the Biomedicum Stem Cell Centre core facility, University of Helsinki, using CRISPR activators^53^. Briefly, fibroblasts were detached as single cells from the culture plates with TrypLE Select (Gibco) and electroporated using the Neon transfection system (Invitrogen) using CRISPRa plasmids^53^. Electroporated fibroblasts were plated on Matrigel-coated plates (growth factor reduced; Corning; 1:200) immediately after the transfections. The medium was changed every other day, and on day 4, the fibroblast medium was changed to a 50:50 ratio of fibroblast medium and stem cell medium (DMEM/F12 with GlutaMAX supplemented with 20% KnockOut Serum Replacement, 0.0915 mM 2-mercaptoethanol, 1x Non-Essential Amino Acids (all from Thermo Fisher Scientific), 6 ng/mL bFGF (Sigma), and 0.25 mM NaB). iPSCs were maintained on Matrigel-coated plates in Essential 8 Medium (E8; Life Technologies). 0.5 mM EDTA was used for lifting cells while passaging, and 5 µM Y-27632 ROCK inhibitor (Selleckchem) was applied into the culture to enhance attachment when thawing.

### Differentiation of iPLCs

The iPLCs differentiation protocol was adapted from Blanchard et al., 2020^18^ with slight modifications. On day 0, iPSCs were dissociated to single cells by Accutase (Thermo Fisher Scientific) and replated on Matrigel-coated plates at a density of 2×10^4^ cells/cm2 in E8 medium with 10 µM Y27632. On day1, the medium was switched to N2B27 medium (1:1 DMEM: F12 and Neurobasal medium supplemented with 1x GlutaMAX, 2% B-27, 1x N2, 50 µM 2-mercaptoethanol and antibiotics (all from Thermo Fisher Scientific)) with 25 ng/ml BMP-4 (PeproTech) and 8 µM CHIR99021(Cayman) for 4 days for mesoderm specification, and the medium was changed every two days. On day 5, medium was switched to N2B27 medium supplemented with 10 ng/ml PDGF-BB and 2 ng/ml TGFβ-3 (both from PeproTech) for two days to induce differentiation toward pericyte-like cells. Then the iPSC-pericytes were maintained in N2B27 medium for an additional two weeks. All the assays were performed on days 21-35 after differentiation started.

### Differentiation of iECs

The iECs differentiation protocol was adapted from K. Wang et al^54^. Initially, the iPSC lines were transduced with ETV2 under the control of a doxycycline-inducible promoter (Tet-On system). The pInducer20-ETV2 plasmid was developed by the Genome Biology Unit, and lentivirus particles were produced by the Biomedicum Virus Core. Both facilities are supported by HiLIFE, the Faculty of Medicine at the University of Helsinki, and Biocenter Finland. On day 0, iPSCs transfected with ETV2 were dissociated into single cells using Accutase and plated onto Matrigel-coated plates at a density of 2.2x104 cells/cm2 in E8 medium supplemented with 10 µM Y27632. On day 1, the culture medium was changed to S1 medium (DMEM/F12 with 1x GlutaMAX, 60 μg/ml L-Ascorbic acid, and antibiotics), and 6 µM CHIR99021 was added for 2 days. On day 3, the medium was transitioned to Stempro medium (Stempro 34 SFM) supplemented with 50 ng/ml VEGF-A, 50 ng/ml bFGF, 10 ng/ml EGF, 10 μM SB431542, and 2 μM doxycycline for another 2 days. On day 5, the medium was further changed to Human Endothelial SFM (Thermo Fisher Scientific) supplemented with 5% Knockout serum replacement (Thermo Fisher Scientific), 10 ng/ml FGFb, 5 ng/ml EGF and 0.5 ng/ml VEGF. Subsequently, the cells can be passaged in this medium and used for experiments for up to 1 week.

### Differentiation of iAstrocytes

The iAstrocytes differentiation protocol was described previously by Oksanen et al., 2017^55^. In short, the medium was switched to neural differentiation medium (NDM, DMEM/F12, and Neurobasal (1:1), 1% B-27 without vitamin A, 0.5x N2, 1x GlutaMAX, and antibiotics) supplemented with dual SMAD inhibitors 10 μM SB431542 (Merck) and 200 nM LDN193189 (Sigma). Then, the medium was changed daily for 12 days until rosette-like structures appeared. Then, the medium was switched to NDM supplemented with 20 ng/ml bFGF for 2 to 3 days to expand rosettes. The areas with rosettes were lifted and cultured in ultra-low attachment plates (Corning) with NDM for 2 days for sphere formation. Next, spheres were cultured in astrocyte differentiation medium (ADM, DMEM/F12, 1x N2, 1x GlutaMAX, 1x MEM-NEAA, antibiotics, and 0.5 IU/mL heparin (Leo Pharma)) supplemented with 10ng/ml bFGF and 10 ng/ml EGF (PeproTech). Spheres were cultured in ADM medium for 5 to 7 months to get pure astroglial cultures. To mature astrocyte progenitor before the assay, spheres were dissociated with Accutase and replated on Matrigel-coated plates in ADM supplemented with 10 ng/ml CNTF (PeproTech) and 10 ng/ml BMP-4 for at least 7 days.

### Differentiation of iNeurons

The iNeurons differentiation protocol was described previously by Kettunen et al., 2023. In short, on day 0, a 60-70% confluent plate of NGN2-iPSCs on was exposed to 2 μg/mL doxycycline in E8 medium. On day 1, the medium was switched to N2 medium supplemented with 2 μg/mL doxycycline and dual SMAD inhibitors (0.1 μM LDN-193189, 10 μM SB-431542B, and 2 μM Xav939). On day 2, the concentration of doxycycline and dual SMAD inhibitors was halved and 5 μg/mL puromycin was added for NPC selection. After removing puromycin and dead cells on day 3, the differentiation continued in N2 medium with full concentration of supplements. By day 4, emerging neurons were plated with or without astrocytes on poly-d-lysine and laminin-coated surfaces. The medium was then changed to Neurobasal supplemented with 1% GlutamMAX, 2% B27, 50 μM nonessential amino acids, 0.3% glucose, and neurotrophic factors (10 ng/mL of GDNF, BDNF and CNTF). Proliferation was halted on day 7 with an overnight treatment of 10 μM floxuridine, and cells matured for 4-6 weeks, with medium changes thrice weekly.

### Immunocytochemistry

Cells were washed with PBS and fixed in 3.7% formaldehyde (Merck) for 20 minutes at room temperature (RT). Then, permeabilized in 0.3% Triton X-100 (Merck) and blocked in 5% normal goat serum (NGS, Merck) in PBS for 1 hour at RT. Primary antibodies were prepared in PBS with 5% NGS and incubated at 4°C overnight. After three washes with PBS, secondary antibodies were applied and incubated for 1 hour at RT. Nuclei staining with 1 µg/ml DAPI (Sigma) was performed for 10 minutes at RT. Primary and secondary antibodies are listed in Table 2. Coverslips were mounted on glass slides with Fluoromount-G™ mounting medium (Thermo Fisher Scientific). Images were visualized by an EVOS microscope (Thermo Fisher Scientific) with 4x and 10x objectives. Brightness and contrast were adjusted in ImageJ software.

**Table 2.** Primary and secondary antibodies used for ICC.

| Target | Dilution | Source | Catalog no. |
| --- | --- | --- | --- |
| Mouse anti- PDGFRβ | 1:100 | R&D Systems | AF385-SP |
| Mouse anti-Chondroitin Sulfate Proteoglycan (NG2) | 1:100 | Merck | MAB2029 |
| Rabbit anti-Actin, α-Smooth Muscle + ACTG2 | 1:300 | Abcam | ab32575 |
| Mouse anti-Oct4 | 1:400 | EMD Millipore | MAB4401 |
| Mouse anti-CD31 | 1:500 | Agilent Dako | M0823 |
| Goat anti-Nanog | 1:100 | R&D Systems | AF1997 |
| Mouse anti-TRA-1-81 | 1:200 | EMD Millipore | MAB4381 |
| Mouse anti-SSEA4 | 1:400 | EMD Millipore | MAB4304 |
| Mouse anti-α-SMA | 1:300 | Sigma | A5228 |
| Mouse anti-B -III-tubulin | 1:1000 | Covance | MMS-435P |
| Mouse anti-AFP | 1:300 | Sigma | A8452 |
| Goat anti-Mouse IgG (H+L), Alexa Fluor<sup>TM</sup> 488 | 1:300 | Thermo Fisher Scientific | A-11001 |
| Goat anti-Rabbit IgG (H+L), Alexa Fluor<sup>TM</sup> 568 | 1:300 | Thermo Fisher Scientific | A-11011 |

### 2D tube formation assay

96-well plates were pre-coated with 50 µl Matrigel and kept in the incubator for 30 minutes to allow Matrigel polymerization. Following that, 3×10^4^ iECs, 1×10^4^ iPLCs (3×10^4^ cells for iPSC-ECs and pericyte-like cells only cultures) were stained with 100nM Calcein AM (Cayman chemical) for 15 minutes at 37°C and seeded the cells on top of the Matrigel layer in a 1:1 mixture of endothelial medium and N2B27 medium. Images were captured every 2 hours using Incucyte S3 (Sartorius) whole well module with 4x magnification, phase contrast, and green channel settings. The analysis of total length, number of segments, and meshes was using 6-hour post-replating images and performed using the Image J Angiogenesis Analyzer module. The images were cropped to exclude shadowed regions, then obtained results were normalized to the analyzing area.

### Permeability assay

iPLCs were seeded on Matrigel-precoated basolateral sides of transwell inserts (0.4 µm pore size, Corning) at a density of 1.5×10^5^/cm^2^. After 24 hours, the apical side of the transwells was coated with Matrigel for 2 hours at 37°C. iECs were then dissociated into single-cell suspensions using Accutase for 5 minutes. Following dissociation, iECs were replated on the Matrigel-coated transwell insert membranes at a density of 4.5×10^5^/cm^2^. Permeability assays were performed 7 days after iECs were replated on the insert membranes. 4 kDa and 70 kDa dextran labeled with Alexa 488, and Texas red fluorophores, respectively (both from Sigma), were mixed with the medium to reach a working concentration of 0.5 mg/ml. Standard curves were generated ranging from 0.5 mg/ml to 160 ng/ml with 5-fold dilutions. 900 µl fresh medium was added to the bottom chamber while 300 µl dextran and medium mixture were applied to the upper chamber. After 1 hour of incubation at 37°C, 100 µl medium from the bottom well was collected and analyzed by FLUOstar Omega spectrometer (BMG Labtech).

### qRT-PCR

Total RNA was extracted from iPSCs, iPLCs, iECs, and iAstrocytes by RNeasy Mini Kits (Qiagen) following the manufacturer’s instructions. RNA concentrations were measured by the SimpliNano Spectrophotometer (Biochrom). RNA was subsequently reverse transcribed to cDNA using the Maxima Reverse Transcriptase kit and supplemented with RiboLock RNase Inhibitor, dNTP Mix, and Random Hexamer Primer (all from Thermo Fisher Scientific). The mRNA expression levels were measured by quantitative RT-PCR using TaqMan assay probes (listed in Table 3.) with Maxima Probe/ROX qPCR Master Mix (Thermo Fisher Scientific) and readout detection by the CFX96 Real-Time PCR System (Bio-Rad). Expression levels were normalized to *GAPDH*.

**Table 3.** primers assay mixes used for mRNA expression studies.

| Gene | Identifier | Source |
| --- | --- | --- |
| PDGFRB | Hs01019589_m1 | TaqMan, Thermo Fisher Scientific |
| DES | Hs00157258_m1 | TaqMan, Thermo Fisher Scientific |
| LAMA2 | Hs00166308_m1 | TaqMan, Thermo Fisher Scientific |
| DLC1 | Hs00183436_m1 | TaqMan, Thermo Fisher Scientific |
| PDE7B | Hs01054008_m1 | TaqMan, Thermo Fisher Scientific |
| CD248 | Hs00535586_s1 | TaqMan, Thermo Fisher Scientific |
| ACTA2 | Hs00426835_g1 | TaqMan, Thermo Fisher Scientific |
| APP | Hs00169098_m1 | TaqMan, Thermo Fisher Scientific |
| LRP1 | Hs00233856_m1 | TaqMan, Thermo Fisher Scientific |
| BACE1 | Hs01121195_m1 | TaqMan, Thermo Fisher Scientific |
| VEGFA | Hs00900055_m1 | TaqMan, Thermo Fisher Scientific |
| ITGA7 | Hs01056475_m1 | TaqMan, Thermo Fisher Scientific |
| COL1A1 | Hs00164004_m1 | TaqMan, Thermo Fisher Scientific |
| EDNRA | Hs03988672_m1 | TaqMan, Thermo Fisher Scientific |
| EDNRB | Hs00240747_m1 | TaqMan, Thermo Fisher Scientific |
| NANOG | Hs02387400_g1 | TaqMan, Thermo Fisher Scientific |
| SOX2 | Hs01053049_s1 | TaqMan, Thermo Fisher Scientific |
| LIN28A | Hs00702808_s1 | TaqMan, Thermo Fisher Scientific |
| CDH5 | Hs00901465_m1 | TaqMan, Thermo Fisher Scientific |
| GAPDH | Hs99999905-m1 | TaqMan, Thermo Fisher Scientific |

**Table 4.**
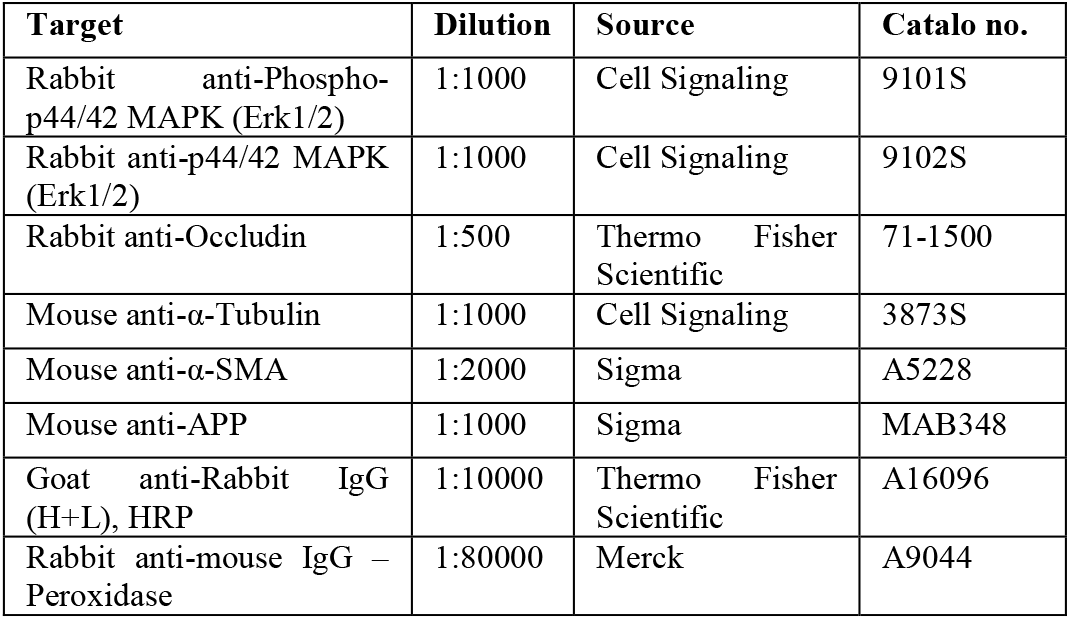
Primary and secondary antibodies used for WB.

### Aβ secretion and uptake

iPLCs were seeded on the 96-well Matrigel-coated plate at a density of 1.5×10^5^/cm^2^. After 6 days of cultivation without medium change, we collected medium from the plates and determined Aβ1-40 and Aβ1-42 levels using the ELISA kit according to the manufacturer’s instructions (R&D Systems). Results were normalized to total protein concentration in the cell lysate (Pierce BCA protein assay, Thermo Fisher Scientific).

Fluorophore-conjugated Aβ1–42 (HiLyte Fluor 488-labeled, AnaSpec) was reconstituted and prepared following the manufacturer’s instructions. Then, 5 µM fluorophore-conjugated Aβ1–42 was prepared in the medium and applied to the culture for 24 hours. After incubation, cultures were washed three times with PBS to remove free residues. 1 µg/ml Hoechst (Thermo Fisher Scientific) was applied to stain the nuclei. The percentage of 488 positive cells was calculated using images taken with the EVOS microscope with 10x objectives.

### iPLCs contraction assay

The assay protocol was adapted from Neuhaus et al., 2017^38^ and Hibbs et al., 2021^39^ with some modifications. Before seeding the cells, 50 µl of the medium was applied into Matrigel-coated impedance plates (E-Plate 16, Agilent) for baseline recording. Then, iPLCs were re-plated on the plate at a density of 6.25×10^4^/cm^2^ and incubated for 30 minutes at RT for cells to attach. The cell index was recorded in 15-minute intervals using xCELLigence® RTCA (Agilent) to ensure the cells reached the confluent stage before the assay, which usually takes 16 to 30 hours. Endothelin-1 (ET-1, Sigma) was added to the culture at a final concentration of 10 nM and the cell index was recorded in 15-second intervals for 2.5 hours and 15-minute intervals for the next 27.5 hours. Since ET-1 was reconstituted in water, an additional 0.1% of water (final concentration) in the medium served as vehicle control. The ‘cell index’ was calculated by software using the cell layer`s resistance to electrical current which reflected the contact area between cells and the well surface. The baseline normalized cell index is generated by normalizing the raw cell index with a mean index of duplicate vehicle controls and the index at the time point when ET-1 is applied to the well. Data analyses were performed with RTCA Software Pro (Agilent).

### Western blot

For tracking the changes of Erk and Akt phosphorylation after ET-1 treatment, 2*105 iPLCs were re-plated on matrigel-coated 4-well plates. One day after seeding, 30nM ET-1 was applied to the well. Cells were collected and lysed after ET-1 treatment for 15, 30, 45, 60, and 90 minutes by radioimmunoprecipitation assay buffer (RIPA) with Halt™ protease inhibitor and Halt™ phosphatase inhibitor cocktails (all from Thermo Fisher Scientific). Protein concentrations were determined by Pierce™ BCA Protein Assay Kit (Thermo Fischer Scientific). 5µg proteins under reducing conditions were separated on 10% SDS-PAGE pre-cast gels (Bio-Rad). Proteins were then blotted onto polyvinylidene difluoride (PVDF) membranes using Trans-Blot Turbo™ Transfer Starter System and Midi PVDF kit (Both are from Biorad). After transfer, membranes were blocked at RT for 1 hour with 5% non-fat dry milk (NFDM) in Tris-Buffered Saline, 0.05% Tween 20 (TBST). Then, the membrane was probed with primary antibodies prepared in 5% bovine serum albumin (BSA) and 0.02% sodium azide in TBST at 4°C overnight. Following three times TBST washes, mouse and rabbit secondary antibodies conjugated with horseradish peroxidase (HRP) were applied and incubated for 1 hour at RT. Proteins were detected and visualized using Pierce™ ECL Western Blotting Substrate or Pierce™ ECL Plus Western Blotting Substrate (from Theromo) and SYGENE Chemi imaging system. Protein relative expression levels were quantified with Image j and normalized to α-tubulin.

To detect α-SMA and APP protein levels, 1*10^6 cells were replated on Matrigel-coated 6-well plates and lysed the following day using RIPA buffer with protease inhibitor. For Occludin and ZO-1 in ECs co-cultured with pericyte-like cells, cells were harvested from transwell inserts using the same lysis buffer. Subsequent procedures were as previously described.

### Cytokines production detection

The Cytometric Bead Array (CBA) with human soluble protein flex sets (BD Biosciences) was used to examine cytokines secreted by iPSC-pericytes. The medium was collected from wells pre-stimulated with 20 ng/ml TNFα and 20 ng/ml IL-1β (both from PeproTech) for 24 hours. Medium was diluted 10 times with assay diluent for TNFα and IL-1β-stimulated samples. 20 µl of samples or standards were incubated with 20 µl beads mixture (75 times dilution) for 1 hour at RT, followed by 2 hours incubation after adding 20 µl detection reagent (75 times dilution). Samples were run on BD Accuri™ C6 Flow Cytometer (BD Biosciences) by detecting around 200 to 300 events for each cytokine. Beads were clustered with 638/660 nm (APC) and 638/780 nm (APC-A750) channels, and cytokines were quantified using 561/585 nm (PE) channel. The data were analyzed with the FCAP array (SoftFlow), and the absolute concentration of secreted cytokines was calculated according to each cytokine’s standard regression curve.

### RNA-sequencing and analysis

iPLCs samples for RNA sequencing (RNA-seq) were harvested on day 21 after differentiation started. The density of cells was evaluated from identical wells ranging from 0.8-1.6×10^6^ cells per well in 6-well plates to ensure line confluency was comparable. Total RNA was extracted using the RNeasy Mini Kit following the manufacturer’s manual. DNase (Thermo Fisher Scientific) was introduced during the RNA isolation procedure to produce DNA-free samples following manufacturer’s manual. RNase inhibitor RiboLock was added after the elusion step to inhibit RNase activity for better RNA preservation. The RNA quantity and quality were analyzed with Agilent 2,100 Bioanalyzer™ (Agilent). The Illumina Stranded Total RNA Prep with Ribo-Zero Plus kit (Illumina) were used to deplete ribosomal RNA and prepare RNA-seq libraries. The libraries were then sequenced with NextSeq500 (Illumina). The libraries preparation and sequencing service were provided by the Biomedicum Functional Genomics Unit (FuGu) at the Helsinki Institute of Life Science and Biocenter Finland at the University of Helsinki. For detection of differentially expressed genes (DEG), raw sequence reads were aligned to human genome GRCh.38 and annotated to gene exons by STAR aligner v2.7.8a^56^ and HTSeq v0.13.5^57^ with GTF v103, respectively. DEG analysis of APPswe pericytes against healthy control samples were performed using the DESeq2 package in R^58^. With the cutoff settings (FDR <0.05 and absolute log2 fold change >1.5), clustering heatmaps were created with R package heatmap 1.0.12 to show DEGs. In addition, using the same cutoff values, pathway enrichment analyses were done by knowledge-based Ingenuity Pathway Analysis (IPA; Qiagen).

### Statistics

Statistical analyses were performed using GraphPad Prism 5.01 and 9.3.1 software (GraphPad Software Inc). Columns were compared with Student’s t-test or one-way ANOVA with Dunnett’s multiple comparison test. Groups with two variables were analyzed by two-way ANOVA with Bonferroni multiple comparison test. Statistical significance was determined with the p-value <0.05. All data in the graphs are shown as mean ± SD.

**Supplementary Figure 1.**
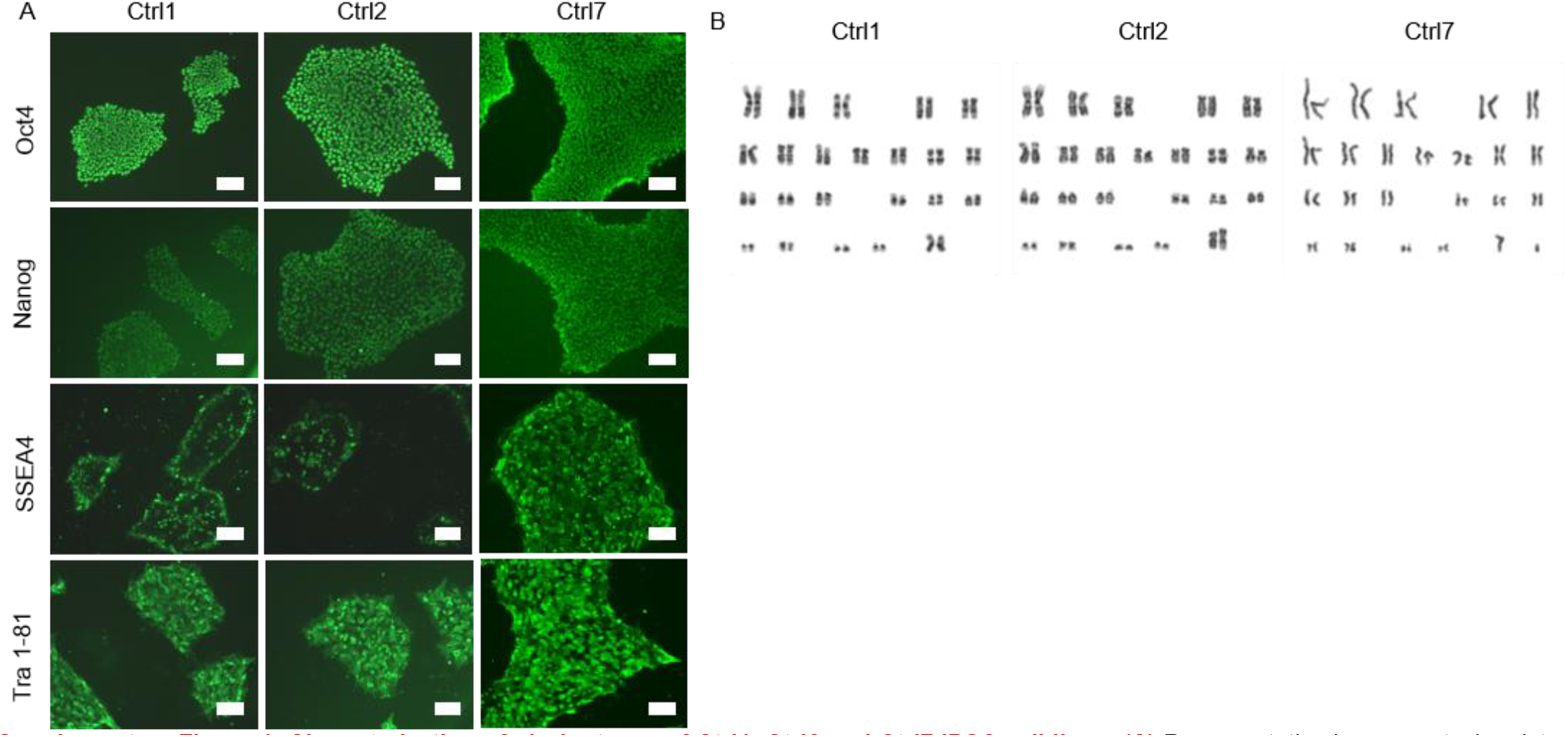
Characterization of pluripotency of Ctrl1, Ctrl2 and Ctrl7 iPSC cell lines. **(A)** Representative immunocytochemistry images of OCT4, NANOG, TRA 1-81 and SSEA4 Ctrl1, Ctrl2 and Ctrl7. Scale bars, 100 µm. **(B)** Representative karyograms from Ctrl1, Ctrl2 and Ctrl7 showing normal euploid karyotypes (46,XX for Ctrl1, Ctrl2 and 46, XY for Ctrl7).

**Supplementary Figure 2.**
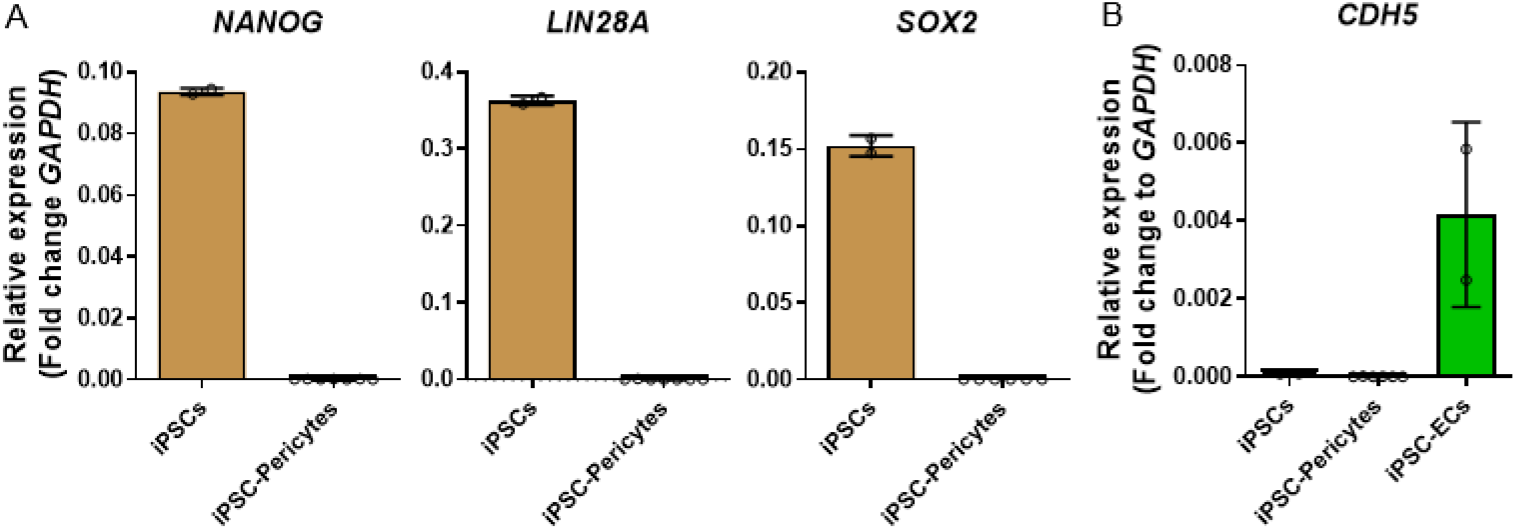
Pluripotency and endothelial markers are not expressed on iPLCs. **(A)** The relative gene expression levels of pluripotency markers *NANOG, LIN28A* and *SOX2* were compared between iPLCs and iPSCs, and were quantified as fold changes to *GAPDH*. The analysis included control iPLCs (n = 6 lines, 2 batches) and iPSCs (n = 2 lines, 2 batches). **(B)** The relative gene expression levels of ECs marker *CDH5* were compared between iECs, iPLCs and iPSCs, and were quantified as fold changes to *GAPDH*. The analysis included control iPLCs (n = 6 lines, 2 batches), iECs (n = 2 lines, 2 batches) and iPSCs (n = 2 lines, 2 batches). The dots represent the average values obtained from batches and technical repeats of each line. The data are presented as mean ± SD. Statistical analysis was performed using one-way ANOVA with Dunnett’s multiple comparison test. The significance levels are denoted as follows: \**p* < 0.05, \*\**p* < 0.01,\*\*\**p* < 0.001 and \*\*\*\**p* < 0.0001.

**Supplementary Figure 3.**
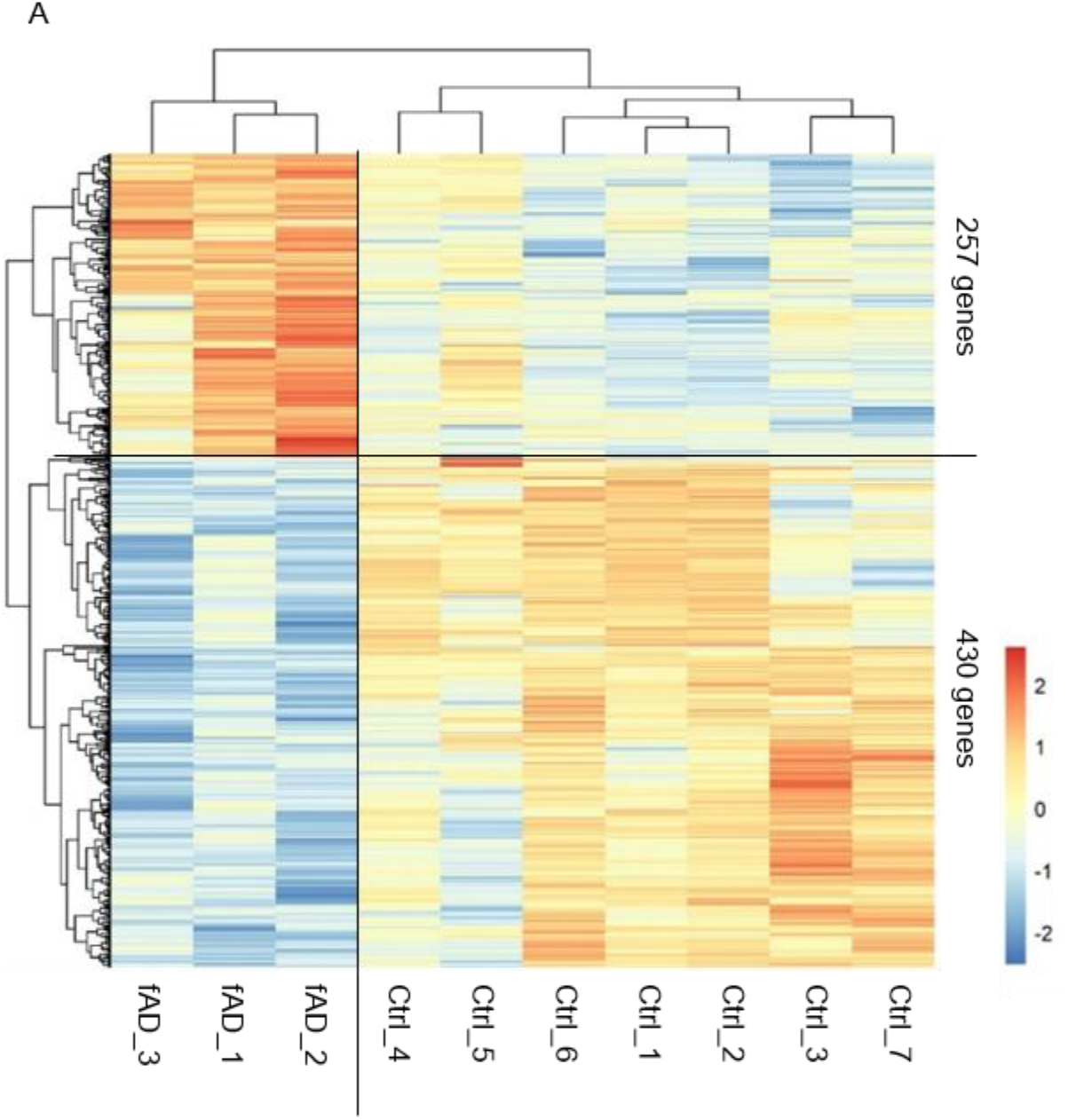
DEGs of APPswe versus control iPLCs. **(A)** Heat map depicting DEGs between control and APPswe iPLCs (cutoffs: Adjusted *p*-value <0.05 and absolute log2 fold change >1.5). The analysis included seven control and three APPswe lines.

